# Mechanistic insights into glucocorticoid-induced ocular hypertension using differences in mouse strain responsiveness

**DOI:** 10.1101/2025.07.02.662542

**Authors:** Pinkal D. Patel, Gaurang C. Patel, J. Cameron Millar, Sherri Feris, Stacy Curry, Eldon E. Geisert, Abbot F. Clark

**Author notes:** Corresponding Author Abbot F. Clark NTERI / IREB536 UNTHSC 3500 Camp Bowie Blvd. Ft. Worth, TX 76107.

## Abstract

Glucocorticoids (GCs) are widely prescribed anti-inflammatory agents. Unfortunately, many people experience negative side-effects associated with long term GC therapy, developing GC-induced ocular hypertension (GC-OHT), which can lead to secondary glaucoma. Approximately, 40% of the treated individuals are susceptible to GC-OHT. Seventy years since this discovery, the molecular mechanisms underlying GC-OHT remain unclear. We previously developed a mouse model of GC-OHT delivering the potent GC dexamethasone (DEX) and observed strain-specific disparities in the development of GC-OHT. We now compare phenotypic and transcriptomic differences between five genetically distinct inbred mouse strains to identify biomarkers of GC susceptibility, and to better understand the molecular mechanisms of GC-OHT. Like humans, mouse strains differ in their ability to develop GC-OHT. Phenotypic characterization revealed that C57BL/6J and C3H/HeJ mice are GC responders and more susceptible to develop GC-OHT. DEX treatment in these strains led to elevated IOP compared to the GC non-responder strains DBA/2J.*Gpnmb^+^*, 129P3/J, and BALB/cJ. Transcriptomic analysis of responder and non-responder mouse strains revealed novel trabecular meshwork (TM) biomarkers of GC-OHT susceptibility involving enrichment of molecular pathways unique to this response. Our study identifies putative mechanisms underlying GC-OHT and provides insight into the pathogenesis of the clinically similar but more prevalent primary open-angle glaucoma.

## Introduction

Glucocorticoids (GCs) are one of the most prescribed classes of anti-inflammatory and immunosuppressive therapies. Currently, an estimated 1-3% of the worldwide population is prescribed glucocorticoid therapy annually (1–4). Since their discovery in the 1940s, synthetic GCs have been a mainstay for treating immunologic and inflammatory disorders, reflecting their increased potencies and specificity highlighting their clinical impact and increased use over the years. Despite their numerous therapeutic benefits, the side-effects associated with chronic GC use collectively contribute to a significant health and economic burden (2, 5).

Elevated intraocular pressure (IOP) is a major ocular side-effect associated with GC therapy, leading to glucocorticoid-induced ocular hypertension (GC-OHT). GC-induced side-effects are observed only in a subset of patients undergoing GC therapy. Approximately, 30-40% of individuals treated with GCs are susceptible to GC-OHT and are GC responders, while the remaining are GC non-responders (6–8). In these susceptible individuals (GC responders), GC-induced insults to the iridocorneal angle tissues causes increased resistance to aqueous humor outflow and elevated IOP, which lead canlead to a secondary form of iatrogenic open-angle glaucoma known as glucocorticoid-induced glaucoma (GIG). The pathology of GIG involves ocular hypertension-induced neuropathic damage to the optic nerve axons, which results in progressive vision loss. Interestingly, the prevalence of GC-OHT is considerably higher (90%) in individuals suffering from primary open angle glaucoma (POAG), the most prevalent form of primary glaucoma (9). Furthermore, first-degree relatives of POAG patients are more likely to be GC responders (∼70-80%) (10, 11). This indicates a potential genetic influence on GC-OHT susceptibility and pathogenesis.

The clinical presentation and disease pathology of GIG closely resembles that of POAG (5, 12, 13). Increased stiffening of the aqueous humor outflow tissues occurs in both diseases. In GIG, this stiffening is associated with GC-induced deposition of excess extracellular matrix (ECM) and cytoskeletal remodeling (14–17) with similar changes occurring in POAG. Given its iatrogenic nature, several animal models of GC-OHT have been developed to study disease mechanisms pertaining to both GIG and POAG (18). We previously developed a reproducible model of GC-OHT in C57BL/6J (B6) mice using periocular delivery of the potent GC dexamethasone (DEX) to elevate IOP (19–21). This and other similar models have been used to study glaucoma pathophysiology. Recently, our group observed a strain-specific variance in GC-OHT susceptibility. Mouse strain background can affect the severity of glaucomatous ocular phenotypes (22). Several studies in mice have shown differences in baseline IOP between genetically distinct inbred mouse strains, indicating a role of genetics in baseline IOP regulation (23, 24).

Clinical evidence shows that severity of GC response varies between individual humans. However, it remains unclear whether GC response rate in mice, across different genetic backgrounds, is homogenous or similar to the heterogenous response reported in humans. To address this knowledge gap, we report our strain survey involving phenotypic and transcriptomic comparisons of five genetically distinct inbred mouse strains: C57BL/6J (B6), C3H/HeJ (C3H), DBA/2J.Gpnmb^+^ (D2.*Gpnmb^+^*), 129P3/J, and BALB/cJ. In agreement with earlier studies (23, 25), we observed differences in baseline IOPs across several genetically distinct mouse strains. Furthermore, we assessed GC-OHT susceptibility by measuring IOP in mice receiving weekly periocular injections of DEX. Among the five strains analyzed, B6 and C3H/HeJ developed significant DEX-induced ocular hypertension (i.e. DEX responder strains). D2.*Gpnmb+*, 129P3/J, and BALB/cJ mice did not develop GC-OHT (i.e. non-responder strains). The GC-OHT response across these five mouse strains is comparable to the ∼30-40% response rate observed in normal humans (7–9, 26). We further performed transcriptional analysis in the TM tissue of responder and non-responder mouse strains to assess mechanisms associated with GC-OHT susceptibility. There was significant downregulation of immune related genes in responder mouse strains compared to non-responder mouse strains. Furthermore, pathway analysis of differentially expressed genes revealed significant enrichment of immune regulatory pathways in responder strains, confirming a stronger GC response in these strains.

## Methods

### Animal Husbandry

Our study examined both male and female mice. Similar findings have been reported in both sexes in terms of GC-OHT response. Genetically distinct parental mouse strains, C57BL/6J, DBA/2J.*Gpnmb^+^*, C3H/HeJ, 129P3/J, and BALB/cJ, utilized in this study were obtained from The Jackson Laboratory (Bar Harbor, ME, USA). All mice were 3-4 months old at the start of experiments. All animal studies and care were performed in compliance with the Association for Research in Vision and Ophthalmology (ARVO) Statement of the Use of Animals in Ophthalmic and Vision Research and the University of North Texas Health Science Center (UNTHSC) Institutional Animal Care and Use Committee (IACUC) regulations (approved protocol: IACUC2023-0027). Mice were housed under controlled temperature (21°C to 26°C) and humidity (40% to 70%), with a 12-hour light/12-hour dark cycle (lights on at 7:00 AM). Food and water were provided ad libitum. The number of animals used in each experiment is indicated in the corresponding figure legends.

### Mouse model of Dexamethasone-induced ocular hypertension

Dexamethasone (DEX) suspension was prepared by mixing 10 mg of micronized DEX (DE121; Spectrum Chemicals) in 1 mL of vehicle (Veh) suspension (Perrone Pharmacy, Fort Worth). Ingredients and preparation of vehicle suspension was described previously (Patel et al, AJP 2017). A uniform suspension with desired DEX particle size was achieved by mixing the suspension along with two stainless steel 5-mm beads (Qiagen, Valencia, CA) in a TissueLyser LT (Qiagen) for 10 min at 50 oscillations/ sec and further rotated overnight at 4 °C until use. Isoflurane anesthetized mice (isoflurane (2.5%); oxygen (0.8 L/min)) were weekly injected bilaterally with 20 μL/eye of either Veh or freshly made DEX (i.e. 200 μg) suspension via the periocular route using a 32-gauge needle attached to a 100 μL volume glass microsyringe (Hamilton Company, Reno, NV, USA). The injection site for the right eye was the inferior fornix and the left eye was the superior fornix of the mouse eye.

### Mouse IOP Measurement

Baseline IOPs were measured on multiple strains of naïve animals as well as on Dexamethasone or vehicle injected animals under isoflurane anesthetized conditions. For isoflurane anesthesia, the mouse was placed in the induction chamber and allowed to inhale a mixture of 2.5% isoflurane and 0.8 liters/min oxygen for 1.5 – 2 mins, or until a deep plane of anesthesia was achieved, indicated by a lack of righting reflex and slowed breathing. Once the desired plane of anesthesia was achieved, the mouse was then immediately removed from the induction chamber and moved to an upright height adjustable stand, where the mouse received a maintenance dose of isoflurane via a nasal cone. IOPs were measured using a TonoLab rebound tonometer (ICare, Finland)(27) stably positioned upright using a metal clamp-stand. This position allowed perpendicular placement of the probe to the central cornea without unwanted movement of animal during measurements, leading to improved accuracy. Each acceptable tonometer reading consisted of six simultaneous button-presses involving six individual readings. The final, sixth button-press, displayed an average of four individual readings while excluding the highest and the lowest reading. We recorded the mean of 7 such IOP readings per eye.

### Mouse TM Isolation

Following four weeks of periocular dexamethasone or equivalent vehicle treatment, mice were euthanized for ocular enucleation and isolation of TM rim tissue. Each mouse was individually euthanized using an IACUC approved protocol involving CO_2_ asphyxiation followed by cervical dislocation. Immediately after a mouse was euthanized, ocular enucleation and TM tissue isolation was performed. Isolation of TM rim tissues was performed in aseptic conditions. Extraocular tissues were first removed with care using extra-fine forceps and scissors. The globe was then punctured ∼1 mm posterior to the limbus using an ocular stab blade. Using the stab-puncture, the globe was hemisected along the posterior limbus and divided into the anterior segment (containing the TM, sclera, ciliary body, iris, lens, and cornea) and the posterior segment (containing the retina). The anterior segment was further cleaned by removing the iris and ciliary body using the extra-fine forceps and scissors. The remaining anterior corneoscleral tissue was folded in semicircle (like a taco). This allowed removal of the cornea using a 4 mm trephine, leaving the TM tissue and the underlying sclera intact. This TM rim tissue was then washed with 1X PBS to remove unwanted cellular or tissue debris. The TM rims from both eyes were then pooled and snap-frozen using liquid nitrogen and stored in a -80 °C freezer for subsequent analysis.

### RNA Sequencing and Pathway Analysis

RNA from mouse TM rim tissues was isolated using RNAeasy Micro kit (Qiagen) protocol as recommended by the manufacturer. Fibrous TM tissues were homogenized by mechanical lysis in the supplied lysis buffer using a TissueLyser LT (Qiagen). Two 5 mm stainless steel mechanical beads were added to the round-bottom 2 mL microcentrifuge tube with frozen TM rim tissues and lysed in TissueLyser at 20 oscillations/sec for 3 cycles of 1 min, separated by 1 min ice incubation after each cycle. RNA quality was determined using a Bioanalyzer (Source). Ribosomal RNA depletion and library preparation was performed using a Zymo Research Library Prep kit (Zymo Research). The resulting library was deep sequenced using the NovaSeq 6K S4PE150 Platform (Microbiome Research Lab at the University of Texas Southwestern (UTSW Dallas)). Reads, QC, mapping, and differential expression analysis was performed using the Qiagen CLC Genomics Workbench. Reads were mapped to *Mus musculus* (house mouse) genome assembly GRCm38. Select differentially expressed (DE) mRNAs were validated at the protein level. Pathway analysis was performed on DE genes using Qiagen’s Ingenuity Pathway Analysis (IPA).

### Western Blots

TM tissues from mouse anterior segments were carefully dissected from enucleated eyes as previously described and were subsequently lysed in Pierce RIPA lysis buffer (ThermoFisher) using sonication (3 cycles of 25 Hz for 3 sec separated by ice incubation). Protein levels in the lysates were determined using Pierce Gold BCA protein estimation kit (ThermoFisher). Equal protein concentrations of lysates were loaded on denaturing 4– 12% gradient polyacrylamide ready-made gels (NuPAGE Bis-Tris gels, Life technologies) for Polyacrylamide gel electrophoresis (PAGE). Proteins from the gels were electrophoretically transferred onto PVDF membranes. Blots were blocked with 10% non-fat milk in TBST solution (1X TBS + 0.1% TWEEN20; Sigma Aldrich) for 2 hours and then incubated for 2 hours or overnight with specific primary antibodies at 4 °C on a rotating shaker at 200 RPM. The membranes were washed thrice with TBST and incubated with corresponding HRP-conjugated secondary antibody for 2 hours. The proteins were then visualized using SuperSignal West Femto Maximum Sensitivity detection reagent (Life technologies). Densitometric analysis was performed on immunoblots using ImageJ (National Institutes of Health; Bethesda; MD).

### Statistical Analysis

Statistical Analysis was performed using GraphPad Prism 10 (GraphPad, San Diego, CA). Data are expressed in means ± SEM. Two-group comparisons were analyzed by Unpaired Student’s t-test. Multiple comparisons were analyzed by Two-way ANOVA followed by the Bonferroni post hoc test. Significance was designated at *P < 0.05, **P < 0.01, and ***P < 0.001.

### Study Approval

This study in animals was approved by the Institutional Animal Care and Use Committee (IACUC) at the University of North Texas Health Science Center at Fort Worth (approved protocol: IACUC2023-0027).

### Data Availability

Values underlying graphed data and reported means from the main text and supplementary files is made available in a Supporting Data Values excel file. RNAseq FASTQ files will be made available via MIAME compliant public database Gene Expression Omnibus (GEO).

## Results

### Mouse genetic background influences susceptibility to GC-OHT

Multiple studies have shown that GC-OHT response in human is heterogenous, and only a subset of the population (30-40%) experience IOP elevation as a side-effect of long-term GC therapy (7–9, 26, 28, 29). Relatives of GC-responders have a greater probability of being GC-responders (10, 11, 26, 29). Glaucoma patients and their first-degree relatives are also more likely to be GC responders (10). Taken together, there appears to be a strong genetic component influencing GC-OHT response in humans (11, 29). Despite this genetic association, studying the GC-OHT response in humans is logistically difficult. Several studies have attempted to identify risk alleles associated with GC-OHT in humans (30–33). In addition, studies utilizing in vitro and ex vivo models have obvious limitations.

Mouse models have been widely used to understand glaucomatous disease pathology and to discover new therapeutic targets. We previously employed the C57BL/6J (B6) mouse model of GC-OHT to study glaucoma-related pathology in the trabecular meshwork and outflow pathway tissues (19–21). The B6 mouse strain is a widely used strain in glaucoma disease modeling, including GC-OHT. Given that all mice share the same genetic background within the strain, variation in baseline IOP is low and DEX treatment leads to significant GC-OHT with near complete penetrance. Previous studies have shown differences in baseline IOPs between genetically distinct mouse strains (23, 25). These inter-strain differences in mouse baseline IOPs are considered analogous to the genetic variation we see between humans. Knowing that only 30-40% of the individuals undergoing long-term GC therapy develop GC-OHT, we asked whether a similar heterogeneity in GC-OHT response is present in mice. To test the genetic association of GC-OHT in mice, we used five genetically distinct inbred mouse strains, which include B6, D2.Gpnmb+, 129P3/J, BALB/cJ, and C3H/HeJ mice. We measured baseline IOP under isoflurane anesthesia in all five strains. Comparison of baseline IOP between the widely used B6 parental strain and other genetically distinct mouse strains showed statistically significant differences (Figure S1). Following measurement of baseline IOP, these strains received weekly periocular-subconjunctival injections of DEX (200 μg/eye) or vehicle, and daytime IOPs were measured weekly in the mornings (Figure 1A). As expected in the B6 strain, DEX injections led to IOP elevation of 4.8 mmHg over the vehicle injected eyes (Figure 1B). The average peak IOP in DEX injected eyes was 23.3 mmHg, whereas in vehicle injected eyes remained at 18.3 mmHg. The C3H/HeJ mouse strain also developed OHT when challenged with DEX. In these mice, DEX injections led to an IOP elevation of 5.0 mmHg, which was comparable to the OHT response we observed in B6 mice. The average peak IOP in DEX injected eyes of C3H/HeJ mice was 18.1 mmHg, whereas in vehicle injected eyes had an average IOP of 13.1 mmHg. To quantify and compare the DEX-induced IOP response in multiple strains, we used the average peak IOP (Figure 1G) and the area under the curve (AUC; Figure 1H). The difference in peak IOP and IOP response (AUC) was determined to be statistically significant (P<0.00001) in the two GC responder strains, B6 and C3H/HeJ (Figure 1G-H). We did not observe statistically significant DEX-induced IOP elevation in D2.*Gpnmb^+^*, 129P3/J, and BALB/cJ strains (Figure 1C-E,G-H), which we refer to as the DEX- or GC non-responder strains.

**Figure 1:**
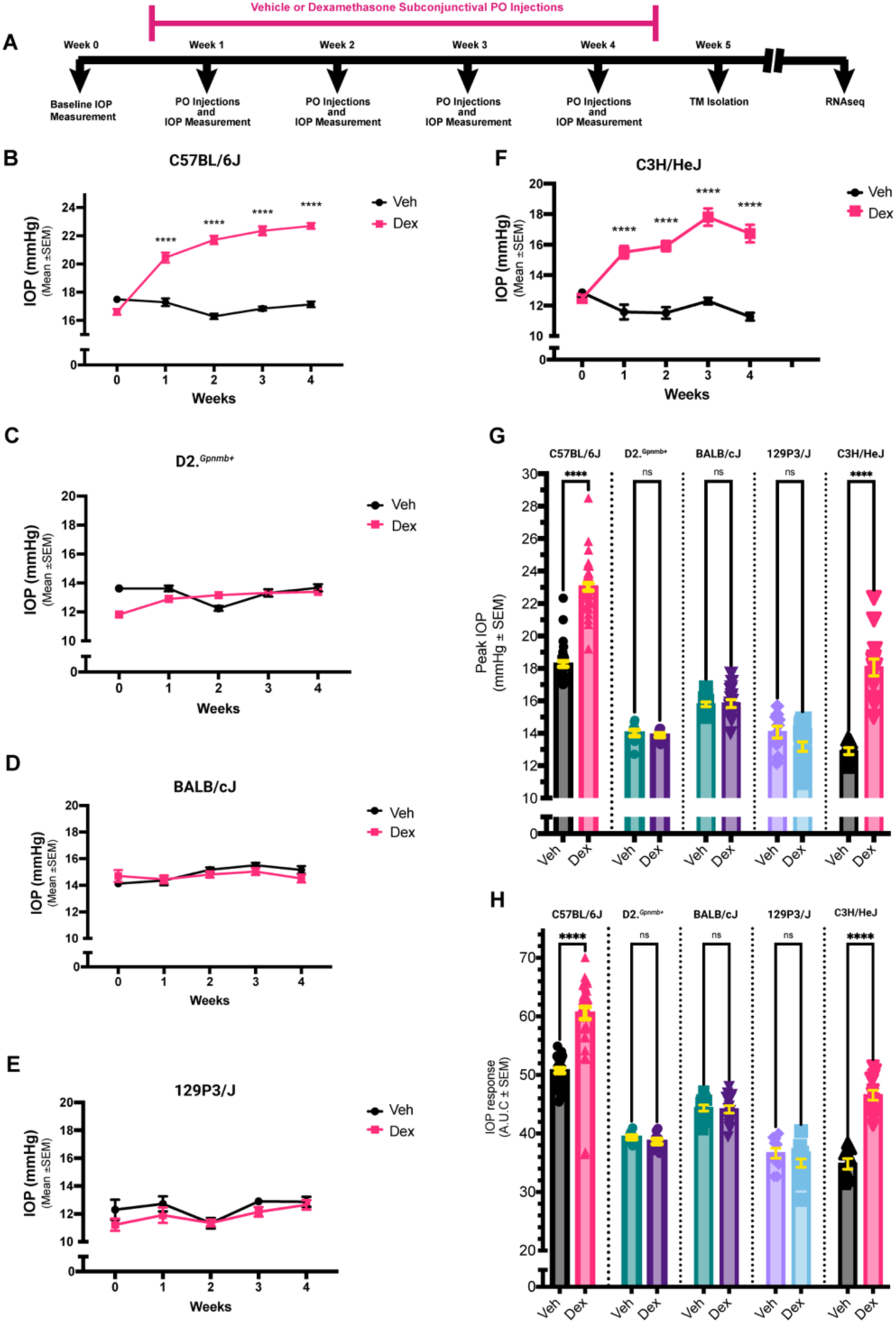
Differential GC-induced ocular hypertension (GC-OHT) response in genetically distinct mouse strains. A) Diagram illustrating the experimental design. Baseline IOPs for each strain were measured. Weekly bilateral injections of Dexamethasone (DEX; 200 μg) or vehicle were administered through the subconjunctival fornix and into the periocular area of the mouse eye to induce GC-OHT. Intraocular pressure (IOP) was measured weekly to determine GC-OHT response. B-F) DEX-mediated effect on IOP in genetically distinct mouse strains. Two-way ANOVA; P<0.0001. G) Comparison of peak IOP attained in mouse strains after DEX or Veh treatment. One-way ANOVA; P<0.0001. H) Analyzing the DIOP response over time (area under curve; AUC) in different mouse strains to assess susceptibility to GC-OHT and identify GC response. One-way ANOVA; P<0.0001.

### Immune regulatory effect of Dexamethasone is stronger in GC responders

Despite clinical association of GCs with the OHT phenotype and glaucomatous pathology, we have yet to identify the mechanisms responsible for GC-induced pathogenic damage to the eye. Given that only 30-40% humans and 40% of mouse strains are susceptible to GC-OHT, we asked which mechanisms make GC responders more susceptible to OHT and glaucomatous damage. In addition, we wonder whether protective mechanisms are at play in the GC non-responders. Our physiological data comparing the IOP phenotype in multiple strains shows that the mouse strains B6 and C3H/HeJ are GC responders, while the other three strains are non-responders. We therefore compared the mouse TM tissue transcriptomes between responder and non-responders to discover the mechanistic differences in gene expression. After IOP measurements, we isolated RNA from TM tissues and performed RNA sequencing (Figure S2). The Venn diagram in Figure 2A shows the numbers of DEX-induced differentially expressed (DE) genes in each mouse strain. A number of differentially expressed mRNAs were unique to the DEX responders. It is important to note that the GC non-responder strain BALB/cJ demonstrated the highest number of differentially expressed genes among the 5 strains analyzed (Figure 2A). This was surprising, given that our phenotypic examination did not reveal a DEX-induced ocular hypertensive phenotype in these mice, so we expected their TM transcriptome to be similar to the other two GC non-responder strains. The experiment was independently repeated in BALB/cJ mice, and DEX-induced changes in IOP were measured via daytime conscious IOP measurement. There was no significant difference in IOP of DEX-injected BALB/cJ mice compared to vehicle injected mice, indicating that the BALB/cJ strain is indeed a GC non-responder strain (Figure S3A). Our analysis further revealed that a vehicle treated BALB/cJ TM sample contained >60% rRNA contamination, which indicates an improper removal of rRNA prior to sequencing. This is perhaps why the BALB/cJ strain showed higher level of differential expression (Figure 2A) because it was highly enriched in several pathways related to RNA metabolism and translation (Figure S3C). The highest-ranking pathways enriched in BALB/cJ were also uncommon to other non-responder strains (Figure S3B), and as a result we chose to analyze the transcriptomic changes in BALB/cJ separately.

**Figure 2:**
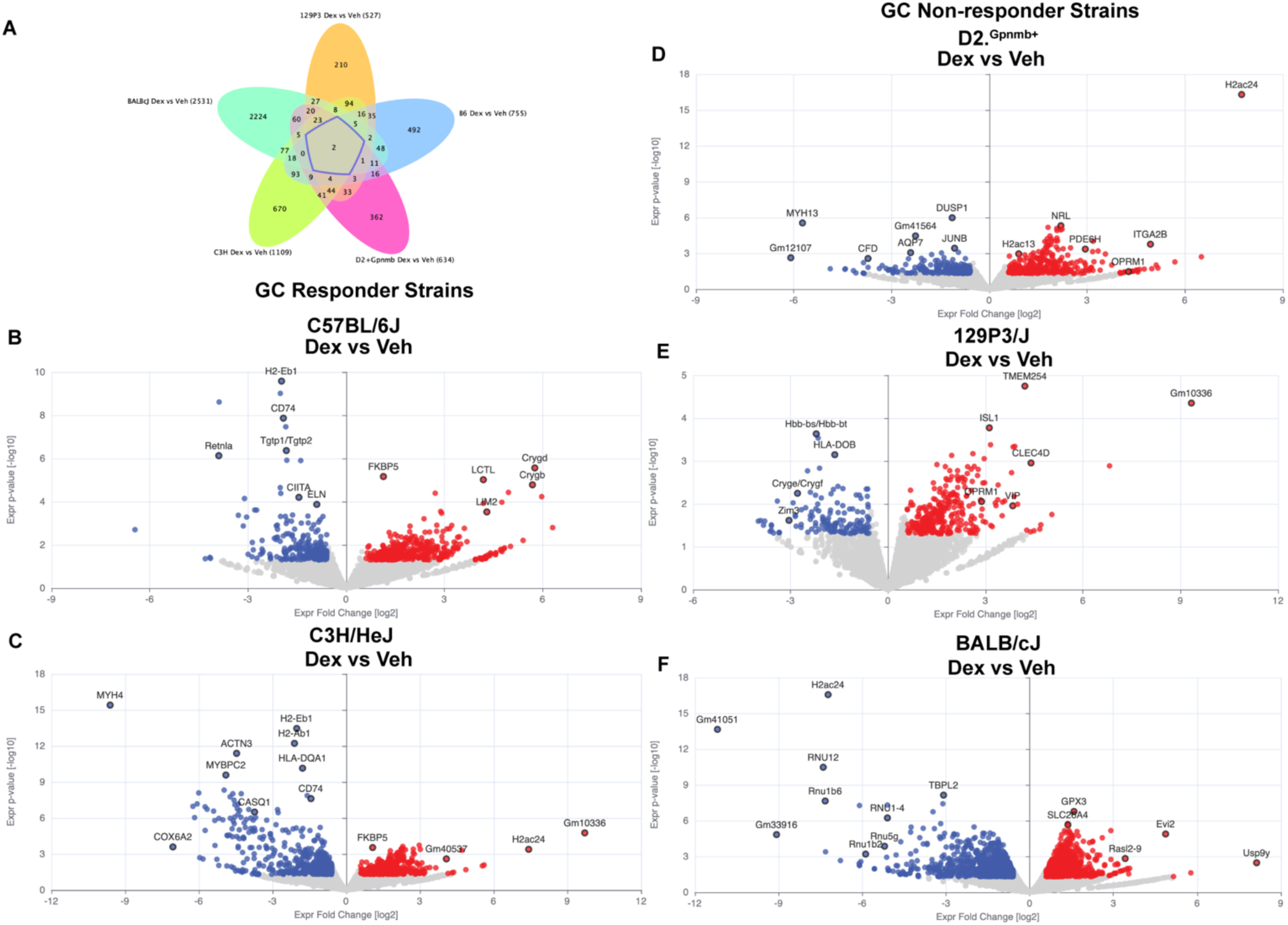
Differential expression of RNAs in GC responder (C57BL/6J and C3H/HeJ) and GC non-responder (D2.*Gpnmb*^+^, 129P3/J, and BALB/cJ) mouse strains. A) Venn diagram showing differentially expressed genes between each strain (P<0.05). B-F) Volcano plot of top 50 most significant differentially expressed mRNAs as a result of DEX treatment in each mouse strain.

As expected, we observed several immune-related genes differentially expressed across all strains (Figure 2B-F). However, the number of these differentially expressed immune-related mRNAs were higher in GC responders when compared to GC non-responders, with significant reduction in overall expression levels. This signified a greater immune regulatory effect in GC responder strains.

### Dexamethasone-induced pathways enriched in GC responders differ from non-responder mouse strains

We then analyzed the DEX-induced differentially expressed mRNAs for each strain using Qiagen’s proprietary Ingenuity Pathway Analysis (IPA) software. All DE mRNAs below the P-value cutoff of 0.05 and beyond fold change cutoff of -/+ 1.5 were considered for this analysis. Figure 3 shows the inter-related pathways enriched in each strain grouped into broader categories and arranged according to the adjusted -log(p-value) for each pathway category. Apart from determining the adjusted P value for each pathway, the IPA also assigns the pathway an activation Z-score determined by the expression pattern of each differentially expressed mRNA belonging to the pathway and expressed in the dataset. We observed highly significant enrichment of pathways related to extracellular matrix organization and immune regulation in the GC responder strains (Figure 3A-B) compared to non-responder strains (Figure 3C-E). We also observed an overall negative Z-score for pathways related to cellular growth, development, and proliferation in the GC responder group compared to the non-responder group. Surprisingly, the GC-responders were also enriched in pathways related to muscle contraction, which was not the case in the GC non-responder strains (Figure 3).

**Figure 3:**
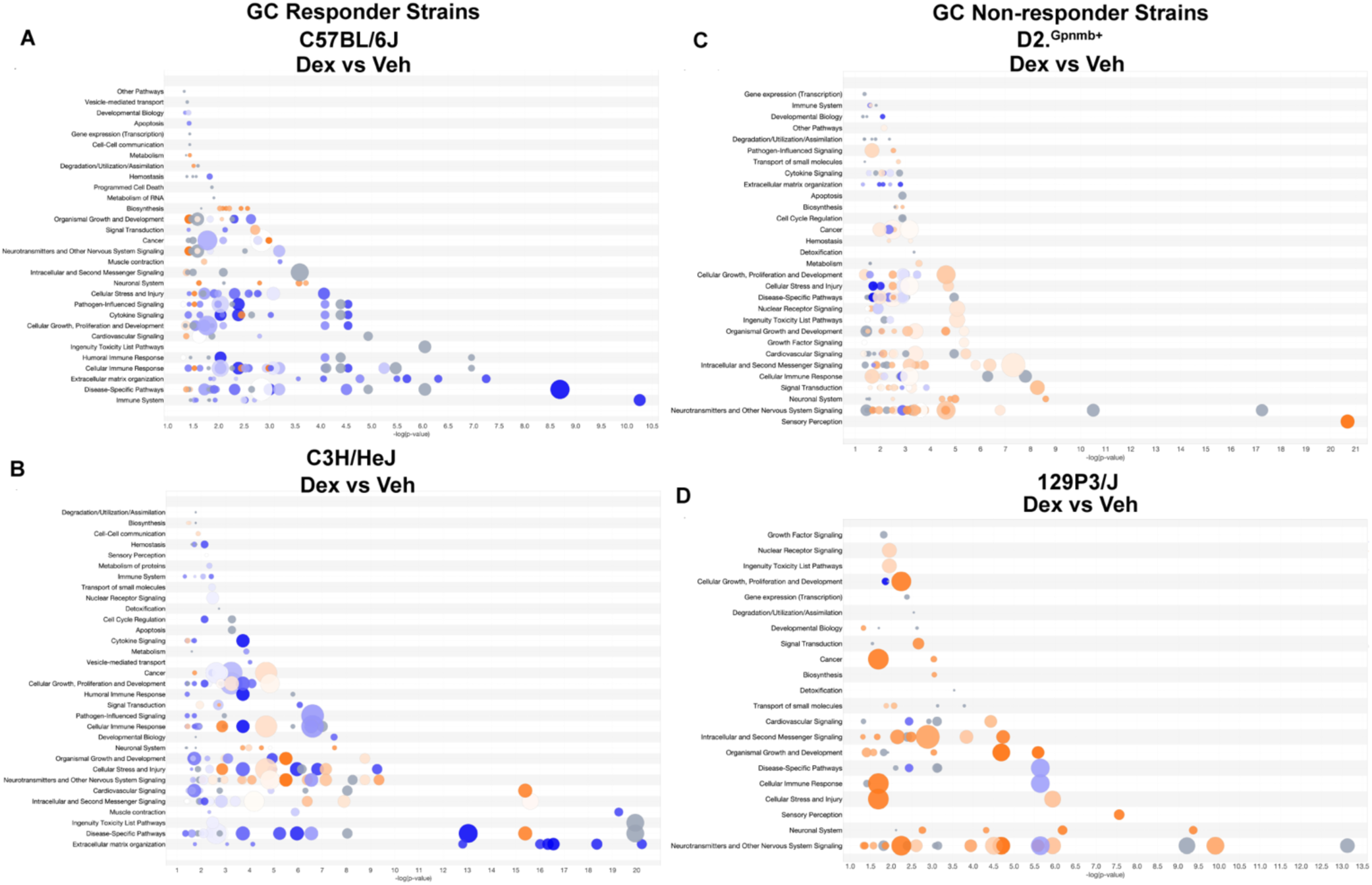
Ingenuity Pathway Analysis (IPA) mediated enrichment of DEX-induced pathways in GC responder strains (A-B) and non-responder (C-D) mouse strains. Related pathways are grouped into a single category.

The GC non-responder strains on the other hand were highly enriched in pathways related to neuronal signaling, second messenger signaling, and neurotransmitter release, indicating robust cell signaling activity at cellular level compared to GC-responders (Figure 3C-E). We observed pathways related to visual phototransduction highly enriched in all the GC non-responders with a high positive activation z-score. We also observed differential expression of photoreceptor (PR) marker gene Rhodopsin (Rho) and retinal ganglion cell (RGC) marker gene POU class IV Homeobox 1 (POU4F1). This uncharacteristic enrichment of vision related pathways and genes in TM samples indicated possible retinal contamination. The small tissue volume of mouse TM and the higher number of retinal cells compared to TM cells increases the likelihood of retinal cells being inadvertently transferred during the dissection process. To address this possibility, we utilized a photoreceptor signature from a large microarray study of the eyes of the BXD strain set (34). The photoreceptor signature was made by taking significant correlates to Rho across the dataset. We then removed from our dataset all genes that highly correlated with photoreceptor marker Rho (1515 genes) and retinal ganglion cell marker Pou4F1 (2001 genes) as previously defined. This included even the genes that were not differentially expressed.

We used the IPA to identify pathways with highest Z-score enriched within the two GC-responder strains C57BL/6J and C3H/HeJ (Figure 4, left). Similarly, we also looked at two of the GC-non responder strains D2.*Gpnmb*^+^ and 129P3/J (Figure 4, right). The heatmaps in Figure 4 show the top ranked pathways enriched in GC-responder and GC non-responder strains determined using the activation Z-scores. Pathways are ranked according to descending order of the adjusted -log(p-values) with the highest ranked pathways at the top. In general, the two responder strains were quite similar to each other in their transcriptomic profiles and distinct from the non-responder strains (Figure 4). The two non-responder strains were also very similar to one another. The GC responder strains (B6 and C3H/HeJ) were enriched in several immune regulatory pathways signifying the potency of immunomodulatory action of GCs in these strains. At a systems level, DEX-mediated immunomodulatory effect in GC responder strains relies on downregulation of cell surface and plasma membrane proteins, which include classes of integrins and immune-related cell surface receptors. Downstream kinase signaling was also reduced in GC responders, which was not observed in the non-responder strains. In the TM, glucocorticoids are also known to induce fibrotic changes involving upregulation of extracellular matrix proteins and dysregulated ECM homeostasis (35, 36). However, in our dataset analyzing long-term effects of GC in mouse strains, we observed antifibrotic changes with downregulation of several ECM genes including collagen and elastin. Apart from the ECM genes, we also observed downregulation of myosin heavy chains and genes regulating intracellular calcium levels, indicating dysregulation of contractile function. In the non-responder strains (Figure 4B, right), we observed DEX-induced upregulation of genes involved in membrane receptor signaling indicating dynamic membrane activity in these cells. To this end, the non-responder strains were also enriched in calcium and potassium signaling pathways with a positive activation Z score. Furthermore, we observed a subdued immunomodulatory effect of GC in non-responder mouse strains as demonstrated by the lack of high-scoring immune-related pathways (Figure 4B).

**Figure 4:**
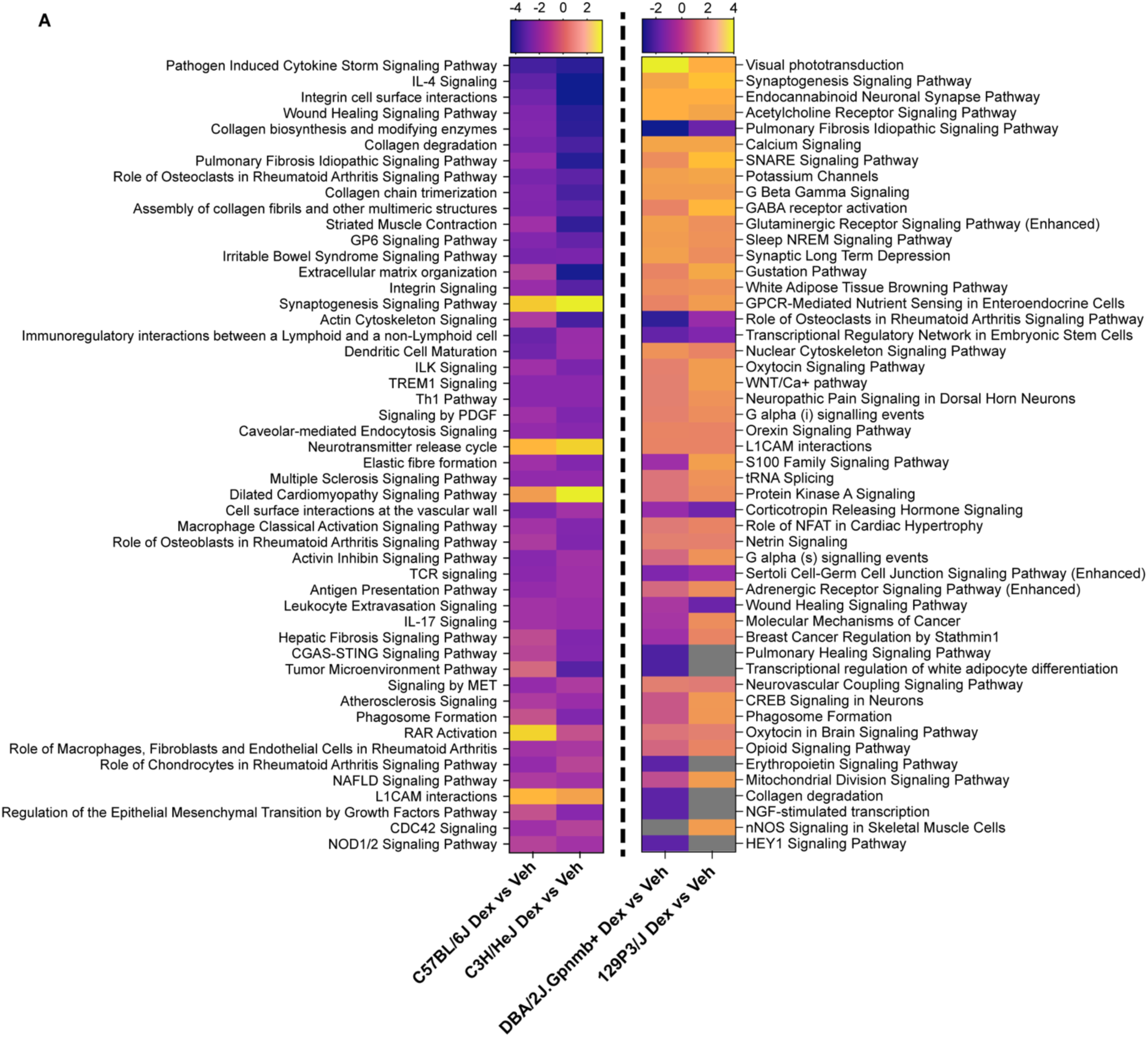
Top DEX-induced canonical pathways enriched based on activation z-score in GC responder (left) and non-responder (right) mouse strains. The z-score indicates a predicted activation or inhibition of a pathway/gene, where a negative z value connotates an overall pathway’s inhibition, and a positive z value connotates an overall pathway’s activation.

### Identification of pathways attributed to susceptibility and protection against GC-OHT

To identify the genes and mechanisms responsible for GC-OHT susceptibility, we grouped the two GC responder strains together (Figure 5A; inset image) and identified 137 differentially expressed genes, which we surmise may be involved in the GC-OHT response. We analyzed these 137 DE genes using IPA and identified a list of pathways common to the C57BL/6J and C3H/HeJ strains (Figure 5A). Likewise, we grouped the two GC non-responder strains, D2.*Gpnmb*^+^ and 129P3/J, and identified 107 DE genes common to these strains (Figure 5B; inset image). Given that GC non-responder strains do not develop DEX-induced IOP elevation, we infer that the DE genes common to these strains may play a role in protection against GC-OHT. We further analyzed these DE genes using IPA and identify DEX-induced pathways prevalent in GC non-responder strains (Figure 5B). Furthermore, IPA identifies the set of differentially regulated genes within our dataset to determine potential the upstream regulators that contributes to the observed expression pattern in each group and predicts the downstream biological function that may be affected. Figure 6 displays the graphical summary highlighting the relationship between differentially expressed genes within the group, their potential upstream regulators, and the related downstream cellular and disease-related functions that are affected in GC responders (Figure 6A) and GC non-responder (Figure 6B) strains. We performed a side-by-side comparison of Z-scores and p-values of GC responder strains and GC non-responder strains using Fisher’s Exact Test. Figure 7 shows the results of these comparative analyses.

**Figure 5.**
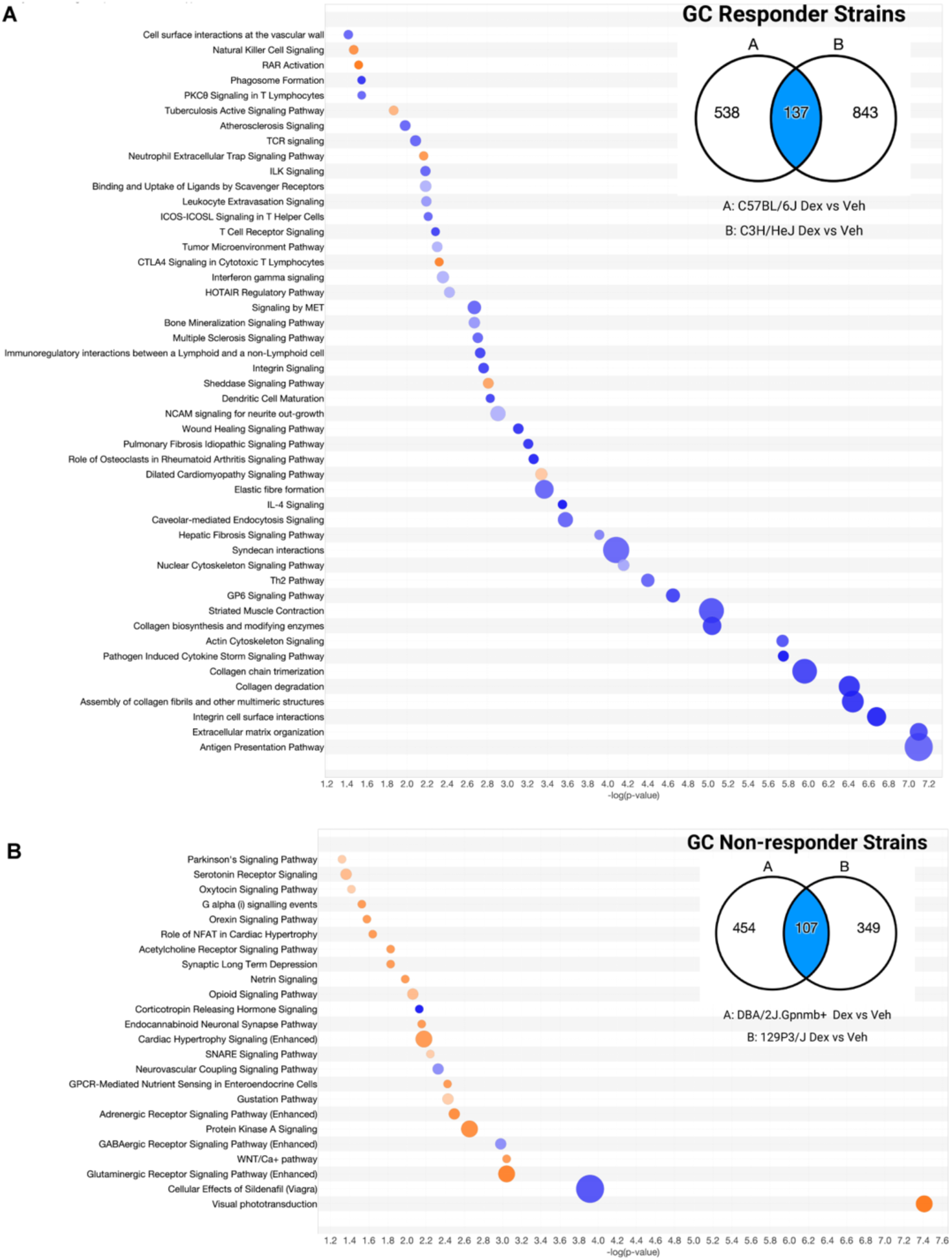
A-B: Canonical pathways associated with GC-OHT susceptibility and protection. A) Pathways contributing to IOP elevation in susceptible GC responder mouse strains. *Inset image*: Venn diagram showing 137 shared DEX-inducible differentially expressed mRNAs (P<0.05) common to GC responder mouse strains. These 137 mRNAs are differentially expressed in response to DEX treatment and are common to the two responder mouse strains (C57BL/6J and C3H/HeJ). *Bottom image*: Diagram showing overlap between the canonical pathways enriched from IPA analysis of 137 DE genes common to responder mouse strains. B) Pathways contributing to protection from IOP elevation in GC non-responder mouse strains. *Top right*: Venn diagram showing 107 shared DEX-inducible differentially expressed mRNAs (P<0.05) common to GC non-responder mouse strains. These 107 mRNAs are differentially expressed in response to DEX treatment and are common to the two non-responder mouse strains (D2.*Gpnmb*^+^ and 129P3/J). *Bottom image:* Diagram showing overlap between the canonical pathways enriched from IPA analysis of 107 DE genes common to non-responder mouse strains.

**Figure 6.**
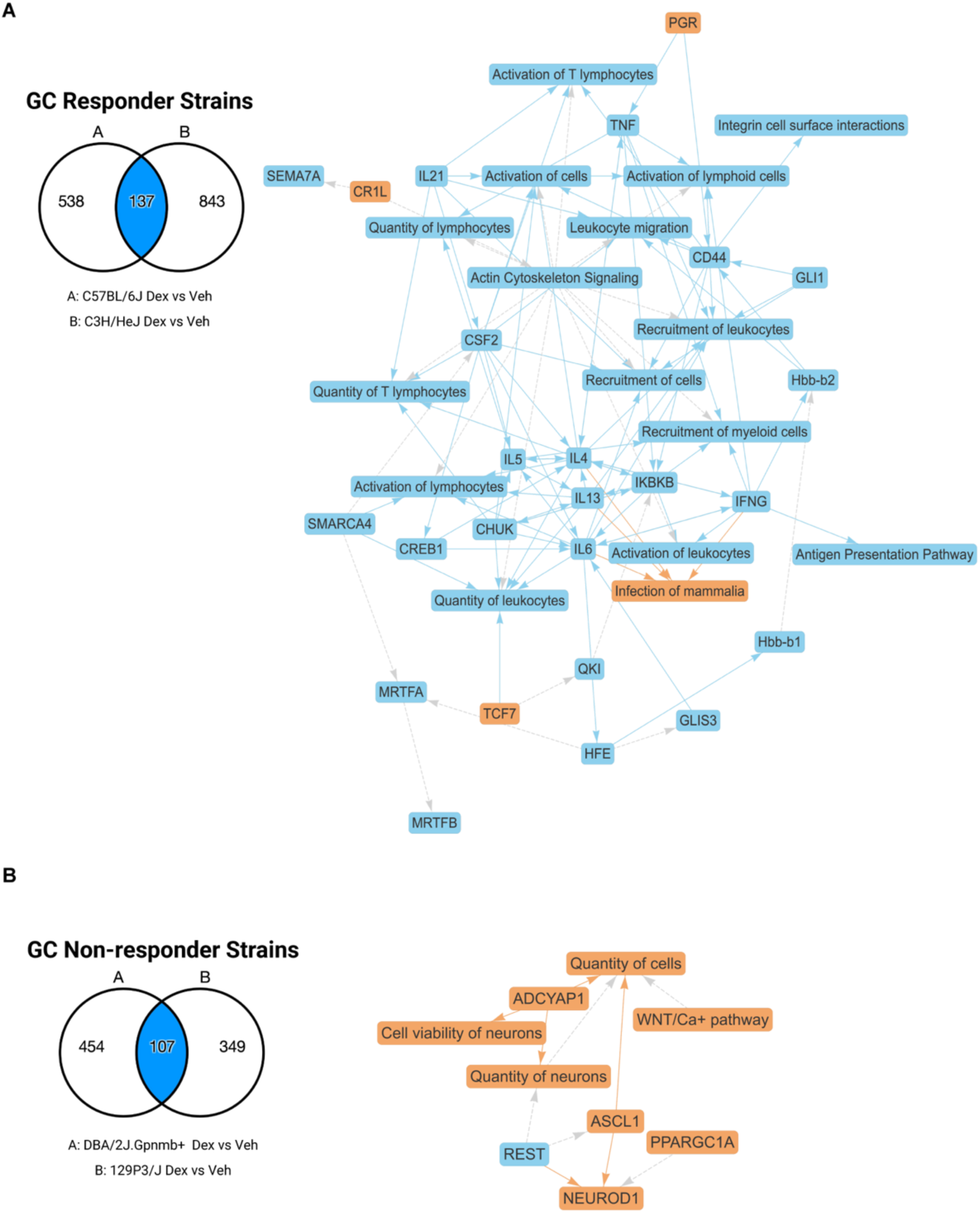
A-B: Graphical summary representing the in-silico IPA analysis performed on genes 137 genes common between two GC-responder strains (A) and 107 genes common between two non-responder strains.

**Figure 7.**
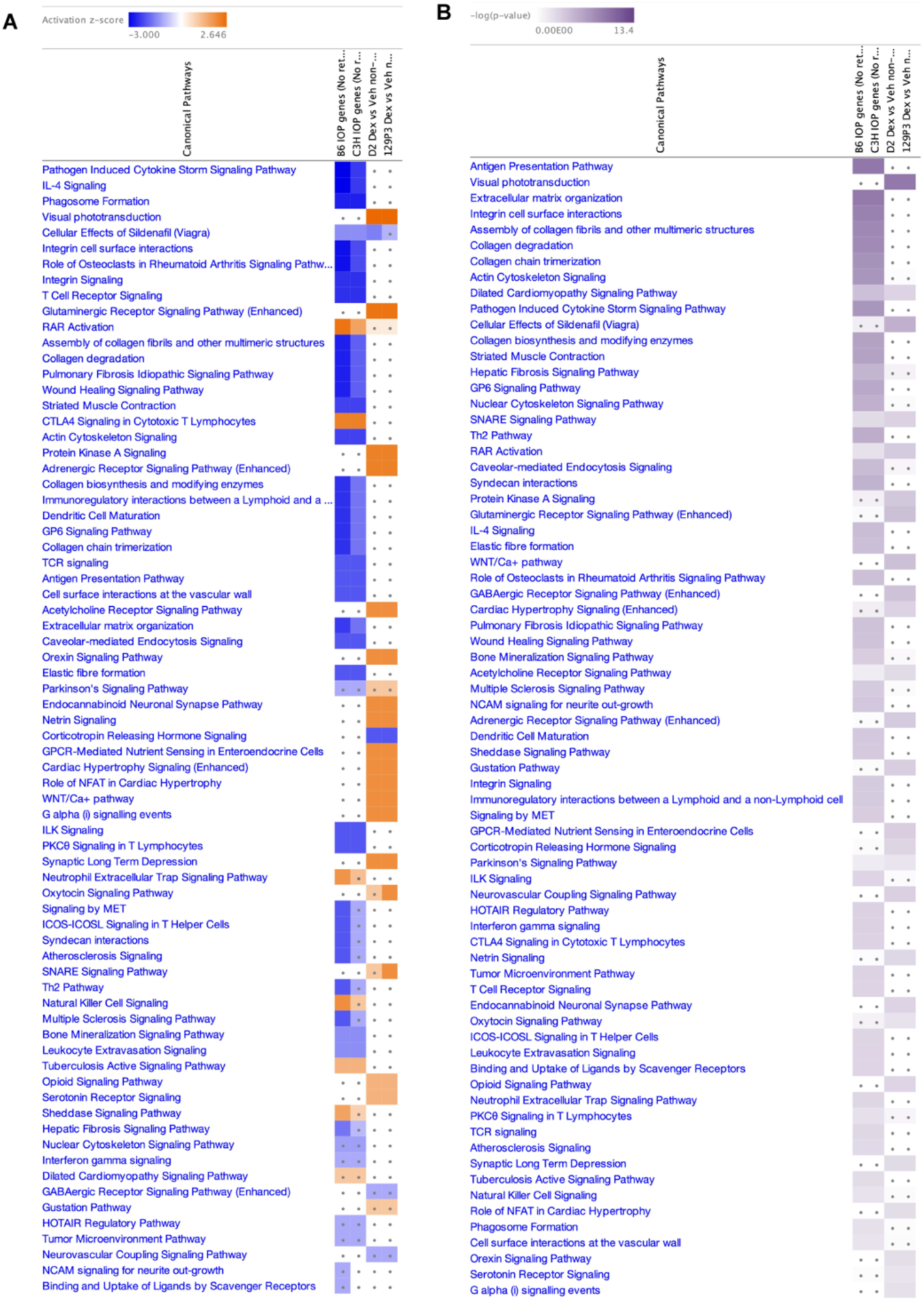
A-B: Comparison of enriched pathways between GC responder and non-responder mouse strains. A-B) Comparison of activation Z-scores (A) and Benjamini-Hochberg corrected p-values (B) for DEX-induced canonical pathways between GC responder mouse strains (C57BL/6J and C3H/HeJ) and GC non-responders mouse strains (D2.*Gpnmb*^+^ and 129P3/J). Pathways are vertically arranged by Z-scores with a dot representing p-value below the insignificant threshold (absolute value 1.3).

### Molecular mechanisms involved in GC-OHT pathology

We used IPA to analyze the DEX-induced DE genes within our dataset and determine possible upstream regulators of DEX-induced IOP elevation. The IPA listed SMARCA4 (Brg1), a catalytic subunit of the SWI/SNF chromatin remodeling complex, as a putative upstream regulator within our dataset, which was predicted to be inhibited (Figure 6A). We analyzed TM tissue lysates from C57BL/6J mice injected with DEX or vehicle for protein levels of SMARCA4 and observed reduced proteins expression in the DEX-treated group (Figure 8A; graph). We also tested protein levels of MRTFA, another upstream regulator identified by IPA (Figure 6A) and known binding partner of SMARCA4. Protein levels of MRTFA were reduced, but the trend was not statistically significant. A previous report in TM cells showed upregulation of SMARCA4 and MRTFA proteins after 5-7 days of DEX treatment (37). This was not observed in our model likely due to difference in treatment timeline. IPA also predicted P38 MAP kinase signaling to be inhibited. We checked for P38 MAPK phosphorylation in DEX-injected TM tissues, which was significantly reduced when compared to vehicle injected controls (Figure 7B). We then used the IPA to unravel the causal relationships associated with our experimental data, which it does by expanding upstream analysis to include regulators that are not directly connected to targets in a dataset but derived from relationships curated from the literature. The causal network analysis (CNA) creates mechanistic hypotheses that may explain the expression changes observed in our dataset. Our CNA results identified Notch signaling as the most prevalent network predicted within our dataset. The CNA result for the master regulator RBPJ/HAT1/NICD/P300 complex (Figure 8C; diagram) was further tested at the protein level. The RBPJ/HAT1/NICD/P300 complex consists of 3 intermediate regulators that control the expression of the several downstream dataset molecules. We observed significant reduction in levels of cleaved Notch Intracellular Domain (NICD) in the TM tissues of B6 mouse injected with DEX compared to that of control (Figure 8C; graph).

**Figure 8:**
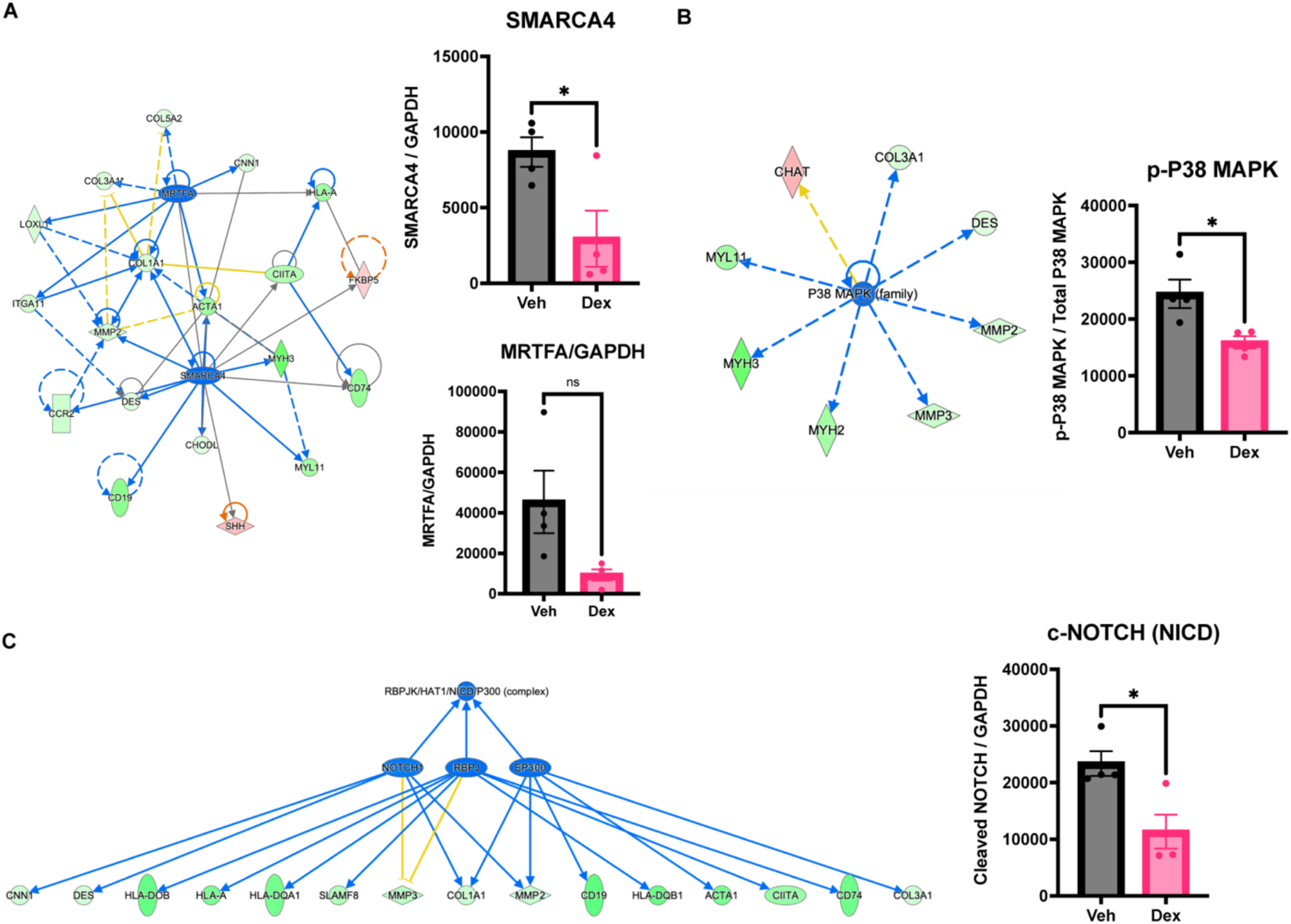
Identification of putative molecular mechanisms involved in GC-OHT response. A) Hub and spoke diagram (left) of the upstream regulator SMARCA4 and MRTFA with its target genes differentially expressed within the dataset. (Right) Quantification of protein levels of SMARCA4 and MRTFA in TM tissues of C57BL/6J mouse treated with DEX or Vehicle; unpaired 2-tailed t-test; *p<0.05. B) Diagram illustrating (left) the P38 MAPK upstream regulator and its target genes. (Right) Phosphorylation level of P38 MAPK against total P38 MAPK in TM tissues of DEX-treated C57BL/6J mice compared to vehicle. unpaired 2-tailed t-test; *p<0.05. C) (Left) Causal network analysis revealed downregulation notch signaling in response to DEX treatment in responders, thereby driving the immunomodulatory effect. (Right) Levels of cleaved notch were reduced in DEX-treated TM rim tissues compared to vehicle; unpaired 2-tailed t-test; *p<0.05.

Apart from predicting upstream master regulators, the IPA also enables identification of putative downstream pathological mechanisms that are at play as a result of the DE genes within our dataset. Hypertrophy is one of the putative downstream pathological mechanisms activated prominently in the responder strains. DEX has previously been associated with increased TM nuclear and cell size but underlying molecular mechanisms have not been thoroughly investigated (38). Figure 8A shows the hypertrophy as a pathological mechanism surrounded by the associated DE genes curated from the literature. Several DE genes from this association were validated at protein level. Protein levels of Elastin (ELN), Junctophilin2 (JPH2), and Triadin (TRDN) were significantly lower in DEX treated TM samples (Figure 8B-D). We also observed reduced protein levels of Collagen1 (COL1) and Myoglobin (MB) in response to DEX treatment, which did not meet the threshold of significance (Figure 8E-F).

## Discussion

GC-OHT is an adverse side-effect of prolonged GC therapy that can potentially lead to glaucomatous neurodegeneration. However, only 30-40% of individuals treated with GCs develop GC-OHT. Such heterogeneity in GC-OHT response has been known for decades, but the underlying mechanisms that make certain individuals susceptible to GC-related pathology remain elusive. Although genetic factors have been associated with GC-OHT response in humans (10, 11, 29), limited numbers of positively identified GC-OHT patients (30–33) make investigating the underlying pathological mechanisms challenging. To overcome this, we decided to use mouse model of GC-OHT previously developed by our group (19–21), which is also now widely adopted to investigate IOP-related pathology in the TM. Using this model, we demonstrate for the first time a heterogeneity in GC-OHT response in genetically distinct mouse strains. Among the five genetically distinct mouse strains tested for susceptibility to GC-OHT, we found two (C57/BL6J and C3H/HeJ) that developed DEX-OHT. This finding, in addition to the distinct baseline IOP observed in these strains, support that genetics may play a role in determining GC-OHT response in mouse, as it has been reported in humans,

This is the first study to compare GC responder and non-responder mouse strains to selectively identify biomarkers and mechanistic pathways relevant to GC-OHT. To compare the genes and pathways that contribute to GC response or GC non-response, we decided to solely focus on candidate genes unique to each dataset. GCs such as DEX are potent anti-inflammatory agents that are involved in immune regulation. There was a strong DEX-mediated change in the transcriptome of GC responder mouse strains compared to non-responders, which involved downregulation of several immune-related genes. This was further confirmed by the comparison of enriched canonical pathways and their respective activity Z-scores between the two groups (Figure 7). The GC responder strains were heavily enriched in immune regulatory pathways with negative Z-scores. This indicated the presence of a DEX regulated immune microenvironment in the outflow pathway of GC responder mouse strains. Although such immune regulation is expected from GC treatment, the strength of the effect was higher only in the two GC responder strains (C57/BL6J and C3H/HeJ), which compelled us to consider that dysregulation in ocular immune environment may be a possible contributor to GC pathology. GC-OHT pathology in the TM, unlike POAG, is reversible in animal and in human. This means that once GC therapy is ceased, the pathology subsides, and IOP returns to normal, which is not the case in POAG. From a functional perspective, TM cells possess several macrophage-like characteristics; they are capable of phagocytosis to maintain the health of the TM by clearing away cellular waste and debris in the aqueous humor, which is vital for proper fluid drainage and regulation of IOP (39–42). Like macrophages, TM cells employ mechanotransduction for sensing and responding to environmental cues like elevated IOP using mechanosensory cation channels (TRPV4, TRPM4, PIEZO) (43–46), cell-cell interacting proteins (cadherins and notch receptors) (47, 48), cell-ECM proteins (integrins) and calcium signaling (49). These cells also possess immune-related markers (TLR4) and secrete immune-regulatory factors (TGFβ) to modulate the tissue microenvironment (50). At systems level, we observe a marked reduction in immune-related surface markers and associated signaling mechanisms in GC-responder strains. This indicates either subtype-specific disruption of immune microenvironment or an overall reduction in immune cells within the outflow pathway. In either case, GC-mediated temporal regulation of immune cells in the TM microenvironment which would hypothetically be reversible upon discontinuation of GC therapy, and subsequent regeneration and reentry of immune cells into the outflow pathway would help revive tissue function. It is yet unclear whether depletion of immune cell populations or change in repertoire of immune cell-subtypes has a role in IOP regulation. Recent reports in multiple systems have identified variety of tissue-resident immune cells playing a role in tissue homeostasis (51, 52). In their recent preprint, Liu et al. demonstrated the presence of long-lived tissue resident macrophages (embryonic origin) and short-lived monocyte-derived macrophages in the anterior chamber that may have a role in maintaining IOP homeostasis (53). This niche of resident immune cells is likely maintained by cues received from the surrounding cells from tissues of the proximal and distal outflow pathways in the form of growth factors and cytokines. If depleted due to stress, these resident-immune cells can be replenished using self-regeneration or via replacement by monocyte derived immune cells (51, 52). This regeneration or replacement of immune cells may contribute to the reversible nature of GC-OHT pathology after discontinuation of GC therapy. A more thorough characterization at the single-cell level is needed to confirm GC-OHT role in altering the resident immune cell repertoire.

From a systems level perspective, we observed that DEX is decreasing immune-related gene expression and regulating immune-associated pathways in GC responders. At cellular level, our in-silico analysis shows several different downstream processes being affected by DEX in the GC responder mouse strains. We see downregulation of ECM proteins and regulatory factors. Cell-cell and cell-ECM signaling is altered with levels of cell-surface receptors and GPCR activity being reduced likely contributing to reduced cell activation. We also observe downregulation of genes responsible for calcium homeostasis and actin cytoskeleton signaling that would affect immune cell mobility within the outflow tissues. Interestingly, we observed DEX-induced downregulation of Junctophilin2 (JPH2) and Triadin (TRDN) at mRNA and protein levels (Figure 9), indicating a possible dysregulation of plasma membrane to endoplasmic reticulum (PM-ER) junctional complexes and intracellular stored calcium release (54–57). Downstream, our in-silico analysis also showed enrichment of actin cytoskeleton and nuclear cytoskeleton signaling pathways both with negative activation Z-scores. These changes may have implication in regulating outflow facility. Actin disrupting agents increase aqueous outflow, and the actin cytoskeleton is rearranged into cross-linked actin networks (CLANs) in TM cells and TM tissues that have been treated with DEX (16, 58–61) as well as TM cells and tissues derived from POAG donor eyes (14).

**Figure 9:**
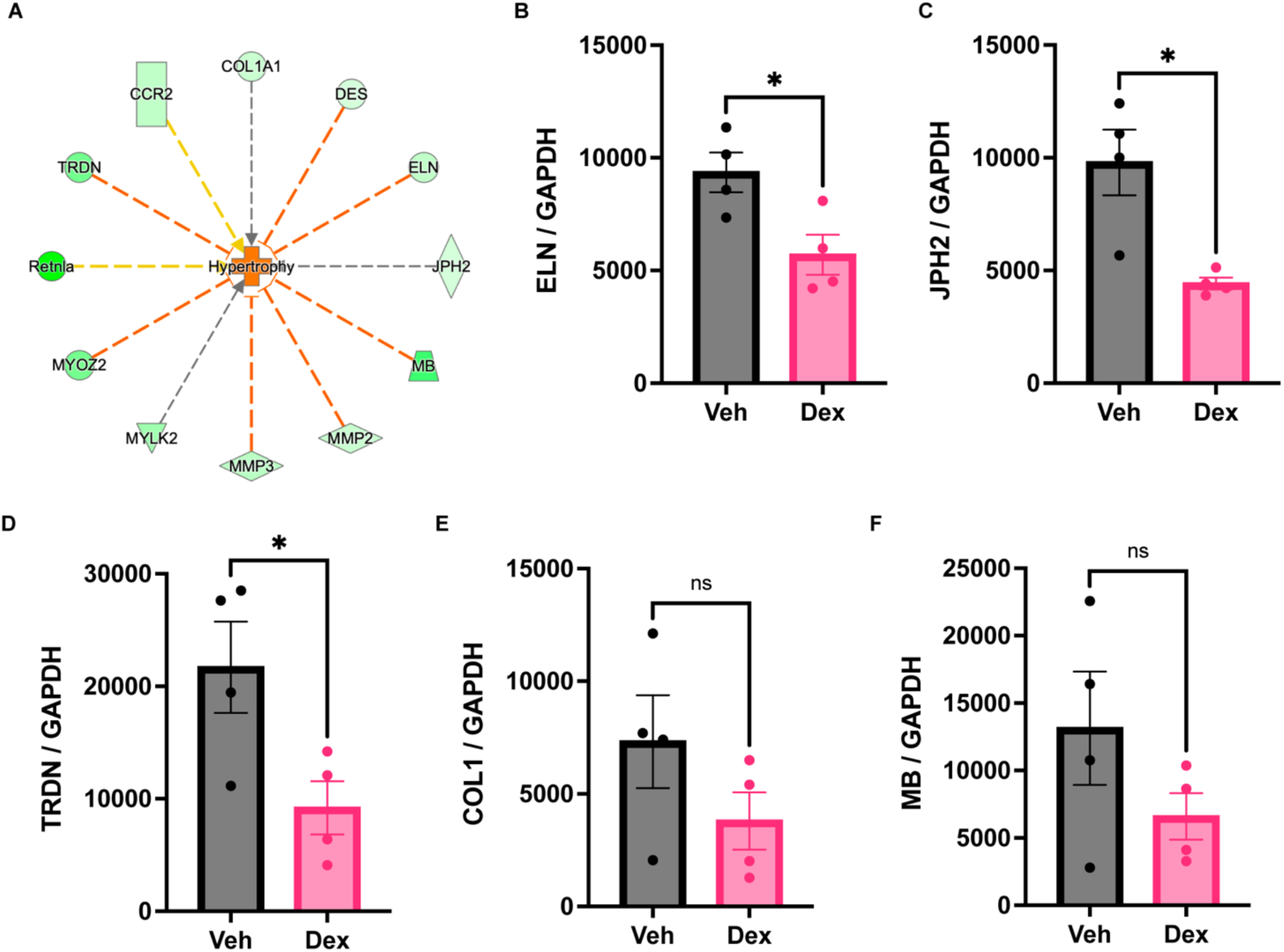
Identification of hypertrophy as disease-associated biological functions involved in GC-OHT response. A) Hub and spoke diagram (left) of hypertrophy as a disease-associated biological function with its target genes differentially expressed within the dataset. B-F) Protein levels of Elastin (ELN), Junctophilin2 (JPH2), Triadin (TRDN), Collagen1 (COL1), and Myoglobin (MB) in TM rims tissues of C57BL/6J mice treated with DEX compared to vehicle treated control; unpaired 2-tailed t-test; *p<0.05.

Multiple processes, such as ECM deposition, focal adhesion, collagen, and TGFβ signaling, have been described to be involved in the pathogenesis of GC-OHT (35, 36). Our GC responder mRNA dataset did not completely align with previous reports. Unlike previous reports, we did not observe significant changes in fibrotic ECM markers except for the levels of collagen and elastin, which were reduced. This difference from previous studies may be due to the prolonged timeline of DEX treatment in our study (four weeks vs 7-14 days). It is possible that this trend could be because of on-going changes in response to GC treatment with TM trying to balance the MMP and TIMP ratios. Apart from the ECM genes, we also tested putative upstream regulators for the DE genes identified in our dataset. SMARCA4, a core subunit of the SWI/SNF chromatin remodeling complex, is essential for development and homeostasis of various organs. We identified SMARCA4 as a strong upstream regulator influencing our dataset of DE genes (Figure 6) in GC responder strains. Given its role in regulating chromatin accessibility and transcription, SMARCA4 may play an important role in orchestrating several processes associated with GC response. A previous study in primary TM cells reported GC-mediated upregulation of SMARCA4 after 5-7 days of DEX treatment, which the authors concluded played a role in downstream actin cytoskeleton remodeling and cell adhesion (37). However, our in-silico analysis using IPA predicted SMARCA4 activity levels to be reduced after 4 weeks of DEX treatment, which we validated and confirmed at the protein level in the TM tissues of C57BL/6J mouse (Figure 8A). Our in-silico analysis also predicted reduced activity of myocardin-related transcription factor (MRTFA), a SMARCA4-interacting transcription factor, as a potential upstream regulator (Figure 8A).

Furthermore, Notch signaling, which involves cleavage of Notch receptor into its intracellular domain (NICD) and the formation of transcription activation complexes, was one of the most highly scored causal networks influencing the expression of gene in our GC responder dataset. We show that NICD protein expression is attenuated in response to GC treatment in C57BL/6J mice. Although the role of notch signaling in IOP homeostasis is unclear, a previous study in human primary TM cells has reported diminished expression of notch signaling molecules with increase in cell-substrate stiffness resembling glaucomatous TM, an effect that also correlated with reduced phagocytosis (47). GC treatment also increased TM substrate stiffness (31). Notch signaling plays a crucial role in tissue homeostasis by regulating activation, differentiation, proliferation, and survival of constituent cells (62). We agree with the sentiments expressed by the previous study (47) that more in-depth analysis of notch signaling is required to understand its potential role in TM pathophysiology.

Although DEX-induced regulatory changes in GC non-responders were diminished, we still observed comparable differential expression of genes. We identified 107 DE genes common between the two non-responder strains (D2.*Gpnmb^+^*and 129P3/J), and these may be responsible for protection against GC-OHT. IPA analysis of this subset of genes had enriched pathways that were different from GC responder strains in function and activity with majority of enriched pathways having positive activation Z-scores. Given the lack of enrichment in immune-related pathways, we asked if there are genetic differences between the glucocorticoid receptors of responder and non-responder mouse strains. To answer this, we mapped the RNAseq reads to mouse genome assembly 38 and performed low-frequency variant detection for identifying variants that may possibly hinder interaction of DEX with its receptor. We used Fisher comparison between GC non-responder strains and GC responder strains to detect variants that were exclusive to GC non-responders. We did not find any variants for the GC binding region but instead found variants in the N-terminal domain (NTD; adj P<0.01). Further analysis revealed that the variant is in the CAG repeat region of the NTD, which is susceptible to repeat expansion and the length of which has been shown to vary between mouse strains. Previous studies in rodents have associated the number of CAG repeats with varying states of GR activation (63–68). It is yet unknown whether the difference in CAG repeat-length has a role in severity of GC-OHT response. We are pursuing this question by employing the BXD recombinant mouse model (69), which represent several inbred recombinant mouse strains derived from crossing the parental strains C57BL/6J (GC responder strain) and DBA/2J (GC non-responder strain). Therefore, we hypothesized that by determining which parental strain contributes the GR gene (Nr3c1) to the recombinant BXD stain, we can predict the likelihood of GC-OHT response in that BXD strain. Our preliminary data involving these recombinant BXD mouse strains that carry the GR gene from either of the two parental strains C57BL/6J or DBA/2J does not appear to correlate with the responder status of the recombinant strain.

### Study Limitations

This study differs in several ways from previous studies investigating GC-induced pathology in the TM. In our prolonged GC-OHT mouse model, we did not observe DEX-induced upregulation of fibrotic mRNA markers in the TM, as previously reported by us and others in human, mouse, and bovine tissues (19, 70–74). This may be due to the protracted timeline of the treatment, type of model, difference in method of tissue isolation and marker detection. Usually the *in vitro* cell-based models and *ex vivo* human and bovine eye perfusion models are used for up to 14 days of DEX treatment. In our *in vivo* mouse model, we extended the treatment time to four weeks. This provides a long-term perspective on GC-induced disease pathology that closely resembles the clinical timeline (8, 28). In fact, our DEX treatment lasted for 4 weeks, at which point we did not observe upregulation of many traditional fibrotic markers within our dataset. This incongruency with previous reports may be because GC-induced elevation of IOP in mouse and human reaches a peak and then plateaus after 2-3 weeks of DEX treatment. This stabilization in IOP may be a form of regulatory mechanism to limit ECM deposition in the TM and to facilitate establishment of a new state of pathological tissue homeostasis. For example, we observed reduced activity of SMARCA4 (Figure 8), which is a chromatin remodeling protein that is associated with fibrosis in TM cells (37). We also acknowledge that there are obvious drawbacks in using mouse eyes for this study. Mouse eyes are very small and complicate the dissection of the TM leading to inadvertent collection of the underlying cells from tissues of SC, outflow vessels, and sclera. Apart from the low tissue yield from mouse eyes, the small number of cells in the TM compared to other ocular tissues like the retina increases the chances of sample contamination with RNA from other cell types. We observed several predefined RGC and PR specific mRNAs differentially expressed in our dataset, which were removed prior to pathway analysis (75). Majority of the reported studies rely on isolated primary TM cells, which excludes the effect of other outflow pathway tissues working synchronously. Although tissues from ex vivo GC perfused human eyes contain multiple tissues/cell-types, they are generically heterogenous and are relatively less fresh (due to the 24-48 delay between time-of-death to collection) than an *in vivo* system like the mouse eye. Therefore, despite the drawbacks, our model provides a comprehensive systems-level image of the putative mechanisms underlying GC-OHT.

### Conclusion

Our results show for the first time a phenotypic difference in the development of GC-OHT among genetically distinct mouse strains. We demonstrate that two mouse strains (C57BL/6J and C3H/HeJ) out of the five analyzed are susceptible to GCs and develop elevated IOP (GC-OHT) in response to prolonged treatment with the potent GC dexamethasone. This is significant because this difference in GC response has been previously reported in humans with a portion of the population being GC responders (∼40%) and the rest considered GC non-responders (∼60%), and we see identical responder rates in mice. We further leverage the mouse strain-specific phenotypic differences in GC response and the power of mouse transcriptomics to gain mechanistic insights into GC-OHT pathology. To our knowledge, this is also the first study to isolate and perform transcriptomic analysis on GC treated mouse outflow tissues. Our transcriptomics evidence suggests that GC responder strains share common pathways and mechanisms that contribute to development of GC-OHT, which are much different from the non-responder strains. We further report a significant inhibition of immune/inflammatory mechanisms in GC responder strains compared to GC non-responder strains. Given the clinical and mechanistic resemblance of GC-OHT to the more prevalent disease of POAG, this work is of broader impact with potential in unravelling molecular mechanisms underlying POAG OHT. This study lays the foundation for future investigations to determine the role of GC-induced immunomodulation in OHT pathology of primary and secondary glaucomas.

---

**Table 1:** List of antibodies used for downstream validation.

| **Antibody Name** | **Catalogue#** | **Dilution** | **Vendor** |
| --- | --- | --- | --- |
| JPH2 Polyclonal Antibody | 40-5300 | 1:500 | ThermoFisher |
| Triadin Monoclonal Antibody (GE 4.90) | MA3-927 | 1:500 | ThermoFisher |
| Myoglobin Polyclonal Antibody | PA5-78396 | 1:1000 | ThermoFisher |
| SMARCA4/ BRG1 Monoclonal Antibody (GT2712) | MA5-31550 | 1:500 | ThermoFisher |
| Elastin Polyclonal Antibody | PA5-76676 | 1:1000 | ThermoFisher |
| LOXL1 Polyclonal Antibody | LOXL-101AP | 1:1000 | ThermoFisher |
| MRTFA/MKL1 Antibody | 21166-1-AP | 1:500 | ThermoFisher |
| P38 MAPK Antibody | 8690P | 1:1000 | Cell Signaling |
| Phospo-P38 MAPK Antibody | 4511P | 1:1000 | Cell Signaling |
| NICD Antibody | 2421S | 1:500 | Cell Signaling |
| COL1 Antibody | NB600-408 | 1:1000 | NovusBio |

