## Supplemental figure S1 for "Mechanistic insights into glucocorticoid-induced ocular hypertension using differences in mouse strain responsiveness": Biorxiv Supplementary Figures 07012025_Figure S1.pdf

A

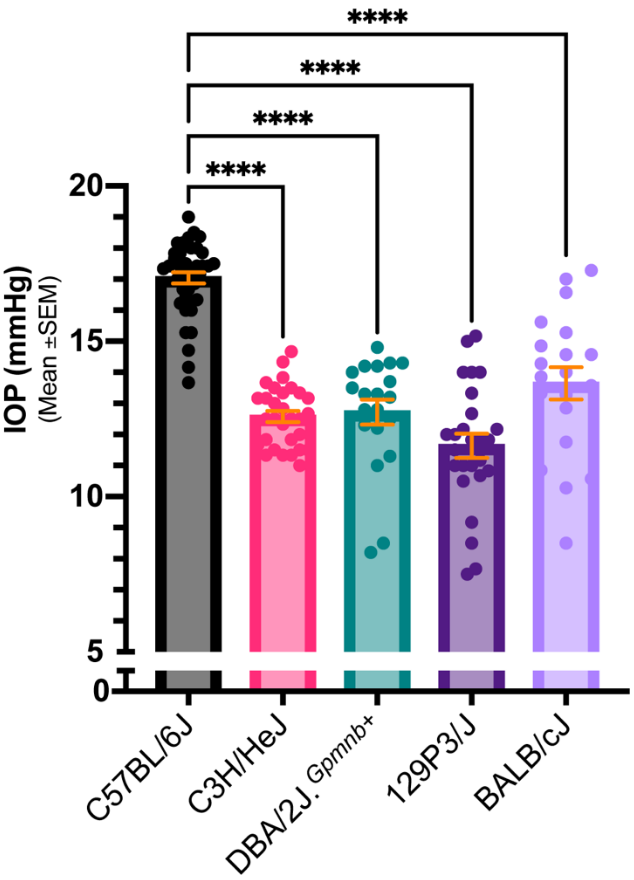


Figure S1: Baseline IOP in genetically distinct mouse strains. Mean IOP (+/- SEM) of each strain is compared with C57BL/6J, a known responder strain. One-Way ANOVA with Bonferroni post hoc analysis for multiple comparisons; P< 0.0001.
