## Supplemental figure S2 for "Mechanistic insights into glucocorticoid-induced ocular hypertension using differences in mouse strain responsiveness": Biorxiv Supplementary Figures 07012025_Figure S2.pdf

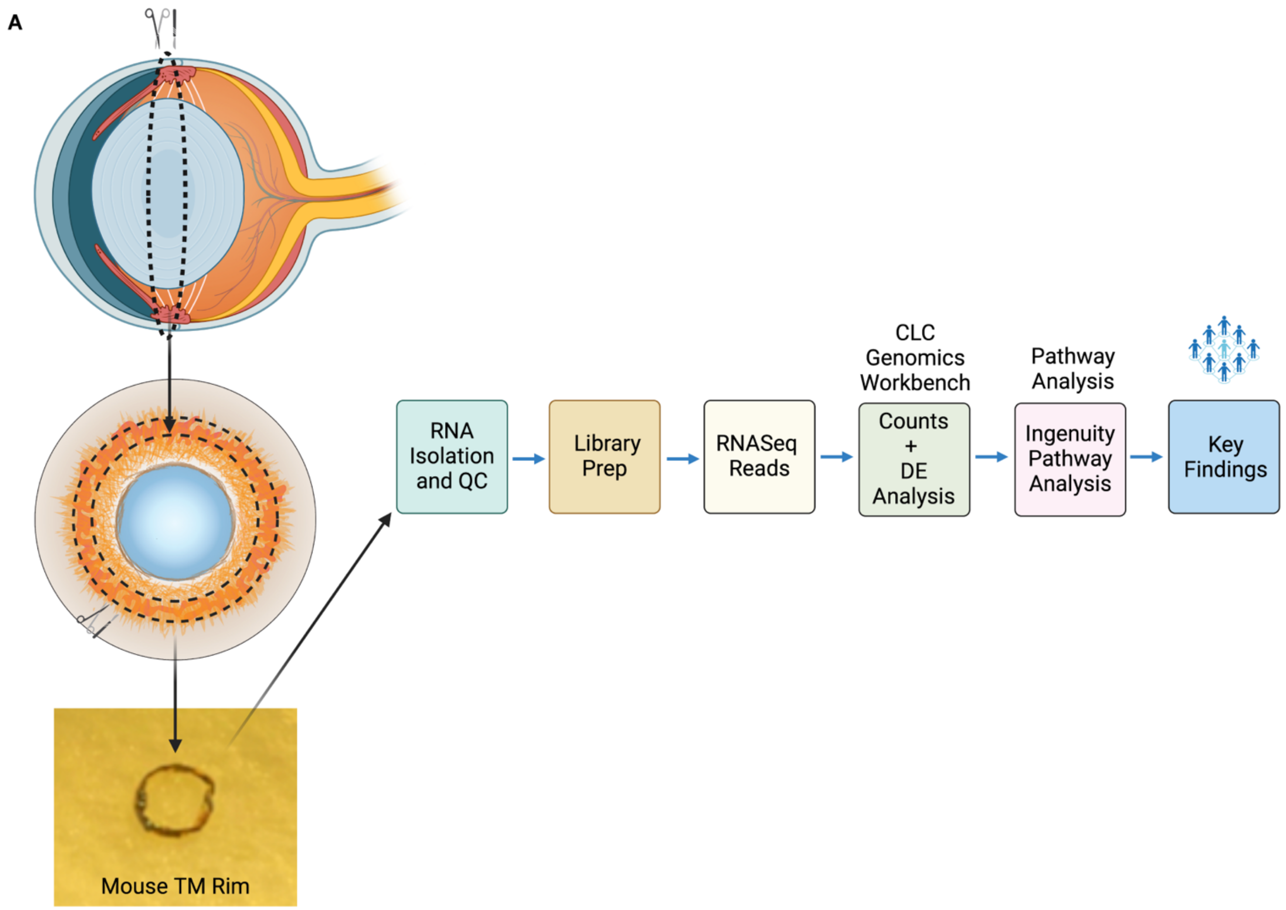


Figure S2: Schematic illustrating TM/Scleral rim tissue isolation and RNAseq analysis pipeline.
