## Supplemental figure S3 for "Mechanistic insights into glucocorticoid-induced ocular hypertension using differences in mouse strain responsiveness": Biorxiv Supplementary Figures 07012025_Figure S3.pdf

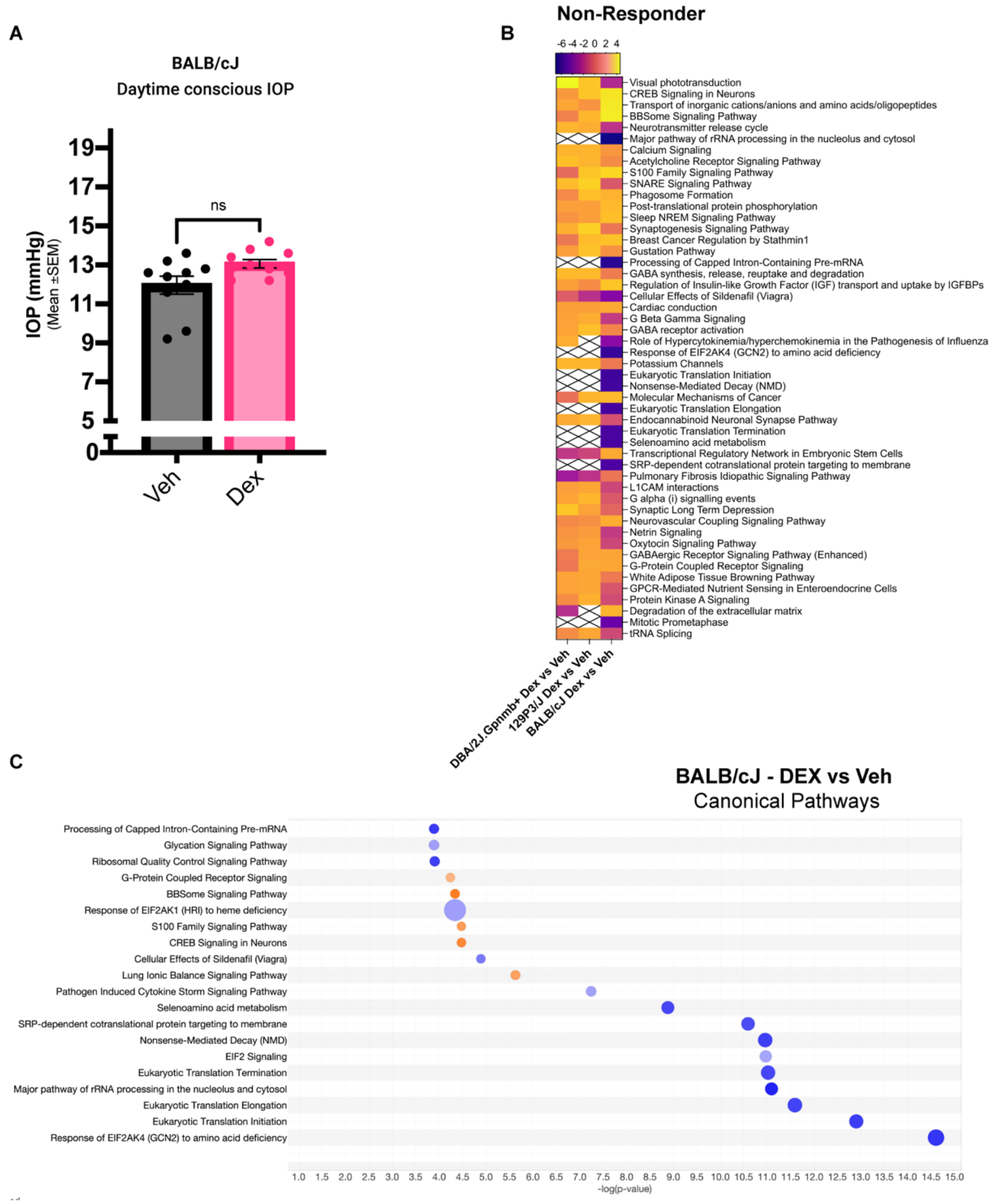


Figure S3: Despite being a phenotypic non-responder, the RNAseq profile of BALB/cJ mouse strain is different from any other mouse strain. This may be perhaps due to possible rRNA contamination. A) Conscious IOP measurement performed in BALB/cJ mouse injected with DEX (10 mg/mL) or equivalent vehicle for 4 weeks. B) Overlap of top DEX-induced enriched pathways in BALB/cJ mouse strain. C) Individual pathways enriched as a result of DEX-treatment in BALB/cJ mouse strain. Red circle denotes pathways related to RNA processing that may likely be due to rRNA contamination.
